# Detection of environmental DNA of the Indo-Pacific humpback dolphins in Hong Kong waters using quantitative PCR

**DOI:** 10.1101/2025.10.30.685690

**Authors:** Robinson O. Kisero, Satsuki Tsuji, Takamitsu Ohigashi, Xiaoqi Lin, Lindsay Porter, Eszter Matrai, Masayuki Ushio

## Abstract

In Hong Kong waters, the Indo-Pacific humpback dolphin (*Sousa chinensis*) is an iconic marine mammal inhabiting shallow coastal regions and plays a vital role in local biodiversity. However, its population is declining, highlighting the need for effective monitoring methods for this species. In this study, we developed a species-specific qPCR assay for detecting *S. chinensis* environmental DNA (eDNA). We designed primers and a TaqMan probe targeting the cytochrome b (*Cytb*) region of mitochondrial DNA. Specificity tests and evaluations of the limits of detection and quantification demonstrated that the primers and probe possess sufficient specificity and sensitivity to detect *S. chinensis* eDNA. Our qPCR method detected *S. chinensis* eDNA in water samples collected from Hong Kong waters where individuals of this species were sighted in close proximity, supporting the field applicability of our method. Analysis of 28 coastal water samples collected in the western and southern Lantau regions of Hong Kong waters in July 2024 demonstrated that *S. chinensis* eDNA could be detected in natural seawater samples. With further enhancements to the eDNA-based survey method, including larger water volumes, broader spatial coverage, and targeting related species and/or taxa such as prey fish, our framework can complement the limitations of traditional surveys, ultimately aiding in the conservation of *S. chinensis* in Hong Kong waters.

## Introduction

Marine biodiversity underpins a diverse array of ecosystem functions, services, and marine resources such as food and energy resources essential for sustainable human society [1].

Unfortunately, biodiversity and its associated functions and services, across both marine and terrestrial ecosystems, are declining due to the adverse impacts of ongoing human activities and climate change [2]. Biodiversity monitoring provides fundamental data to understand the temporal trends of biodiversity, and it is necessary to avoid further declines in the biodiversity, and consequently, ecosystem functions and services. Traditional monitoring methods, such as direct visual censuses, have long been employed to assess the presence/absence and abundance of marine species such as fish and cetaceans [3–5].

However, these approaches are often labour-intensive and costly, necessitating taxa-specific expertise, which limits their scalability and frequency.

Environmental DNA (eDNA), broadly defined as DNA released into the environment by organisms, has emerged as a powerful, complementary tool for biodiversity monitoring, which enables detection of macro-organisms in various ecosystems, including aquatic and terrestrial environments [6–11]. This genetic material carries species-specific information, allowing for the detection of species from a single environmental sample through molecular techniques. For example, conventional or quantitative PCR is usually a single-species detection approach and has been used to detect freshwater amphibians, marine fish, and marine mammals [12–14]. eDNA metabarcoding, which amplifies target genetic regions of a particular taxon and distinguish multiple species by sequencing, has been used to detect multiple species belonging to marine fish, mammals, and freshwater invertebrates [7,15–19]. Recently, eDNA shotgun metagenome has been shown to be effective to study multiple taxa such as vertebrates, invertebrates, microbes and viruses at once [8]. These studies have demonstrated the effectiveness of eDNA techniques as a non-invasive tool for detecting cryptic, elusive, or rare taxa, and the insights gained from eDNA-based biodiversity monitoring can help establish conservation strategies.

In Hong Kong waters, the Indo-Pacific humpback dolphin (*Sousa chinensis*), popularly known as the Chinese white dolphin, is an iconic marine mammal living in shallow coastal regions (Fig. 1a). It is distributed along the coastline of the eastern Indian and western Pacific Oceans [20], and in the South China Sea, the Pearl River estuary and its adjacent waters including Hong Kong waters are home to about 2,500 individuals of *S. chinensis* [21]. In Hong Kong waters, *S. chinensis* is one of resident cetacean species alongside the Indo-Pacific finless porpoises (*Neophocaena phocaenoides*). They serve as an important indicator of the biodiversity and ecological status of the marine environment because apex predators are often sensitive to environmental pollutions and disturbance [22], and because cetaceans can play critical roles in maintaining ecosystem functions such as carbon and nutrient cycling [23,24]. However, due to various factors such as vessel operations (e.g., underwater noise), construction noises and habitat loss, its population size is decreasing [25,26], and currently, *S. chinensis* is categorized as “Vulnerable” by the IUCN [27].

**Figure 1.**
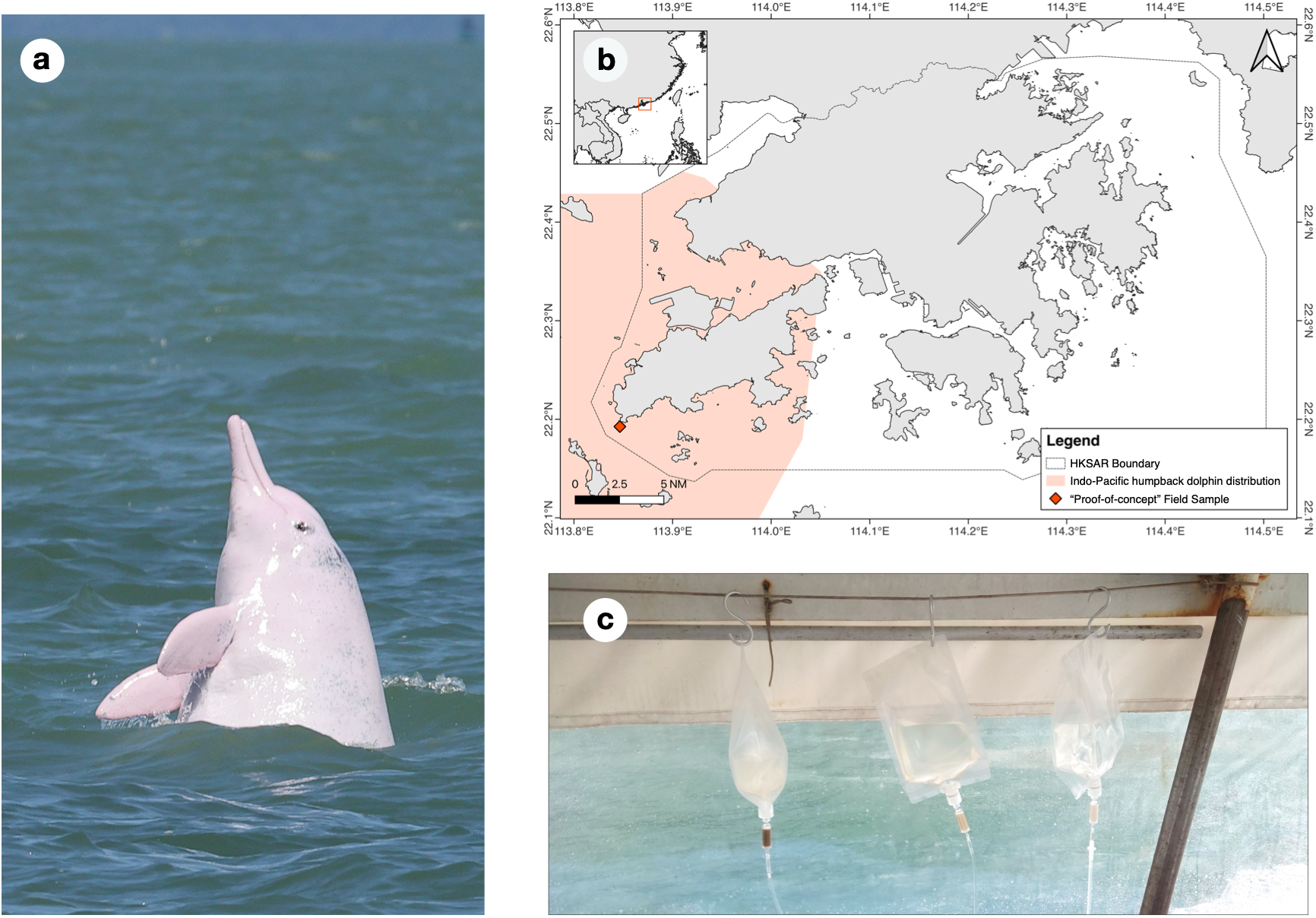
Indo-Pacific humpback dolphin in Hong Kong waters. **(a)** Indo-Pacific humpback dolphin *(Sousa chinensis* ), or Chinese white dolphin, in Hong Kong waters (Photo taken by E. Matrai). **(b)** The distribution of S. *chinensis* around the Lantau island in Hong Kong waters. The red diamond marks the location where we sighted S. *chinensis* and collected the "proof-of-concept" field samples for qPCR analysis. **(c)** Gravity filtration of seawater on a boat during the field survey (Photo taken by M. Ushio). The photos are published under the CC BY 4.0 license.

To conserve its population and even reverse the declining trend, frequent and broad spatial-scale monitoring is necessary to understand population dynamics and the underlying mechanisms. The Hong Kong government has used several survey techniques to monitor *S. chinensis* population and inform with conservation strategies. Some of these techniques include: the photo identification (Photo-ID) survey [28] and systematic long-term line transects survey method [27] using vessels. However, such surveys are often expensive, rely on good weather windows, are labour-intensive, and require the cetacean species under study to surface frequently and remain at the surface long enough. Acoustic-based methods are also used for detecting *S. chinensis* [29], but audiograms can vary among individuals of different age groups, necessitating careful parameter settings for accurate detection. To complement these existing techniques, developing an alternative, potentially scalable approach is highly valuable for more effective and frequent monitoring of Indo-Pacific humpback dolphins in Hong Kong waters.

To complement traditional monitoring methods, eDNA analysis offers a promising alternative for detecting *S. chinensis* in Hong Kong waters, providing several advantages, including relatively low on-site labor requirements, the ability to detect detailed genetic information [30,31], and the ability to detect cetaceans even when they are submerged or have left the immediate survey area. Recently, cetacean eDNA has been detected by several eDNA metabarcoding studies [17,32,33], but the use of quantitative PCR (qPCR) has been particularly beneficial in studies targeting a single or few species, as it allows for sensitive detection and quantification of eDNA which can be a rough index of species abundance or biomass [14,34]. For example, Hashimoto et al. (2024) developed species-specific qPCR primers for East Asian finless porpoises (*Neophocaena asiaeorientalis*) and detected finless porpoise eDNA even in the absence of direct sightings [13]. More recently, Sun et al. (2025) developed qPCR primers for detecting eDNA of *S. chinensis* [35], but the performance of these primers, including the limit of detection, the limit of quantification, and specificity, was not empirically evaluated.

Given this background, the overall aim of this study is to develop a species-specific qPCR assay for *S. chinensis* eDNA and demonstrate its applicability for monitoring the species in Hong Kong waters. Specifically, our objectives are twofold. First, we aim to design primers and a probe specific to *S. chinensis*. We will employ the TaqMan qPCR method [36], as it enables more specific detection of eDNA compared to non-probe based methods (e.g., the SYBR Green method). We then empirically evaluate the performance of the developed method, specifically assessing the limit of detection (LOD) and limit of quantification (LOQ) using synthesised *S. chinensis* DNA fragments, as well as its specificity using synthesised DNA of closely related, potentially co-occurring cetacean species. Second, we apply the developed qPCR method to real seawater samples collected from the coastal region of Hong Kong, focusing on the western and the southern Lantau Island area, a major habitat for *S. chinensis* [26] (Fig. 1b), to demonstrate the potential of eDNA-based detection of *S. chinensis* in Hong Kong waters.

## Methods

### Ethic statement

This study used tissue samples of *S. chinensis* from stranded individuals found in fields to design and validate specific qPCR primers and probe for *S. chinensis*. All tissue samples were obtained with the necessary permits from the Agriculture, Fisheries and Conservation Department (AFCD) of Hong Kong, and Ocean Park Hong Kong (OPHK) and Ocean Park Conservation Foundation Hong Kong (OPCFHK). Seawater samples from Hong Kong waters were collected outside the marine protected areas. No direct interaction with or sampling from live *S. chinensis* occurred during this research.

### Identification of the potentially suitable region and inspection of intraspecific variations using tissue-extracted DNA

To identify a suitable region for our target species, we downloaded the full mitochondrial genomes of 71 cetacean species, 34 non-cetacean species, and 8 common mammals (e.g., humans, dogs, cats) from the NCBI Reference Sequence Database (https://www.ncbi.nlm.nih.gov/refseq/; accessed 11 July 2023; Table S1). This set of mitochondrial genomes is the same as that used to design cetacean-specific eDNA metabarcoding primers in Ushio et al. (2025) [17]. We aligned all sequences using the online MAFFT alignment tool [37], visually inspected them, and identified a potentially suitable region in the cytochrome b (*Cytb*) gene.

To assess intraspecific variations in the *Cytb* region, we collected 29 tissue samples of *S. chinensis* from stranded individuals stored by OPHK and OPCFHK (Table S2). The samples included different age groups (from neonates to adults) and sexes. We extracted genomic DNA using the DNeasy Blood & Tissue Kit (Qiagen, Hilden, Germany) following the manufacturer’s protocol. The extracted DNA was amplified using manually designed primers targeting 913 bp of the *Cytb* region: 5′-GACACCTCAACTGCTTTCTCATC-3′ (forward) and 5′-GGCTGTTGGTATTAGCACTAGGA-3′ (reverse). PCR was performed using a 20 µL reaction volume containing 10 µL of Platinum SuperFi II PCR Master Mix (Thermo Fisher Scientific, Waltham, MA, USA), 2 µL of each 5 µM F/R primer, 6 µL of ultrapure water, and 2 µL of DNA template. The thermal cycle profile after an initial 30 sec denaturation at 98°C was as follows (30 cycles): denaturation at 98°C for 10 sec; annealing at 60°C for 10 sec; and extension at 72°C for 30 sec, with a final extension at 72°C for 5 min. The PCR products were purified using AMPure XP (Beckman Coulter, Brea, CA, USA) and sent for Sanger sequencing. The sequences were aligned using an online MAFFT alignment tool, and we visually inspected them and designed the primers and probe in the region where there is no intraspecific variation.

### Primer and probe design

We designed qPCR primers and a probe specific to *S. chinensis* by considering the following criteria for primer and probe design. We did not design a TaqMan probe in the existing marker region [35] because our *in silico* investigation indicated that the primers developed by Sun et al. (2025) [35], for example, SCDloop-F/R, had zero mismatches (100% identity) with several non-target cetacean species (e.g., *Pseudorca crassidens*, *Tursiops truncatus*, and *Delphinus delphis*; some of these species distributions potentially overlap with *S. chinensis*). Because complete sequence identity in the primer binding sites makes it highly likely that non-target cetacean DNA would be co-amplified, we excluded that region from our analysis and instead designed a new primer–probe set targeting the *Cytb* region. The primers were broadly following general primer-design guidelines: they are between 19 and 28 bp in length, with a GC content of about 40–60%, *T_m_* values of forward and reverse primers at 60±1°C, an amplicon length of 95–170 bp, and a T at the 3′-end avoided. The TaqMan probe was designed to be 25 to 28 bp long, has a GC content of 50% and *T_m_* values of 67–70°C, and is positioned at least 10 bp apart from each primer region. By visually inspecting the *Cytb* sequences of the 71 cetacean species and Sanger sequence data of *S. chinensis*, we designed the following primers and probe in the *Cytb* region (Table 1): qCWD-cytb-F (5′-TCGCTTTCCACTTTATCCTTCCC-3′), qCWD-cytb-R (5′-GATGTCTTTGATTGTATAATAGGGGTGA-3′), and qCWD-cytb-P (5′-AGCAGCCGTTCACCTGCTATTCCTACAT-3′; modified the 5′-end by FAM and the 3′-end by BHQ1).

**Table 1.**
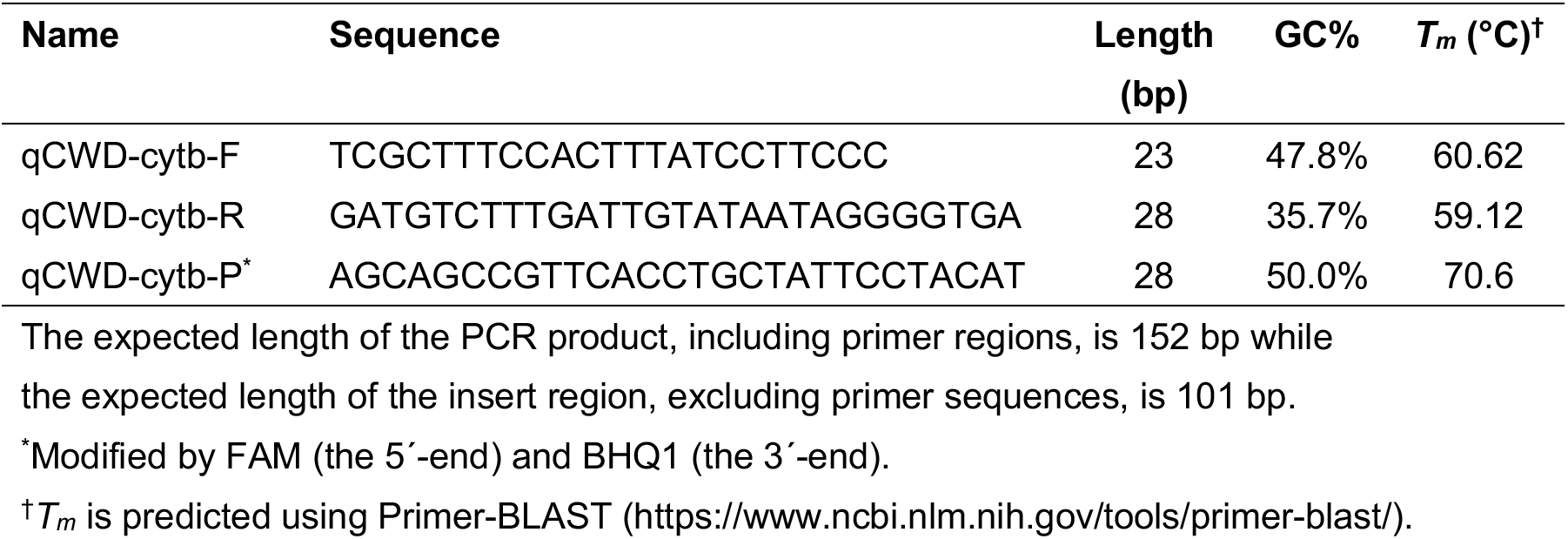
Indo-Pacific humpback dolphin-specific qPCR primers and probe.

The specificity of the primers was first checked using Primer-BLAST (results are summarized in Table S3). We found that one registered sequence of Atlantic humpback dolphin (*Sousa teuszii*) had only one mismatch with the primers, but their distribution does not overlap with that of *S. chinensis* [20]. The common bottlenose dolphin (*Tursiops truncatus*) had three to five mismatches (most registered sequences had four mismatches) with the primers, and one registered sequence of the long-beaked common dolphin (*Delphinus capensis*) had three mismatches. Three and four mismatches were found in many of registered sequences of the Weddell seal (*Leptonychotes weddellii*) and the striped dolphin (*Stenella coeruleoalba*), respectively. Among them, the three cetacean species, *T. truncatus*, *D. capensis*, and *S. coeruleoalba* were occasionally found in Hong Kong waters, so we empirically tested whether the qPCR primers and probe amplify their DNA using synthesised DNA (see the section “*Specificity, LOD, and LOQ of the qPCR primers and probe*” and Results). All other marine species have at least five mismatches with the primers according to Primer-BLAST (for details, see https://github.com/ong8181/qpcr-cwd), so we did not test the amplification of other marine species.

### Specificity, LOD, and LOQ of the qPCR primers and probe

We obtained synthesised DNA of the target region (152 bp; Table S4) of *S. chinensis* from Integrated DNA Technologies (Coralville, Iowa, USA) as standard DNA for our qPCR experiment, and tested the amplification of this standard DNA. Briefly, serially diluted standard DNAs were prepared using the synthesised DNA fragment (1.0 – 10^9^ copies/µl). qPCR was performed in triplicate for each standard DNA concentration on a QuantStudio Flex 6 (Thermo Fisher Scientific) using a 20 µL reaction volume containing 10 µL of TaqMan Environmental Master Mix 2.0 (Thermo Fisher Scientific), 2 µL of primer-probe mix (each 9 µM F/R primer and 1.25 µM probe), 0.2 µl of AmpErase Uracil N-glycosylase (UNG) (Thermo Fisher Scientific), 5.8 µL of ultrapure water, and 2 µL of standard DNA. The thermal cycle profile after an initial activation of UNG at 50°C for 2 min and initial denaturation at 95°C for 10 min was as follows (55 cycles): denaturation at 95°C for 15 sec; annealing and extension at 60°C for 1 min. The fluorescence intensity was measured after the extension step.

We also tested the amplification of the following DNAs to check the specificity of the primers and probe: genomic DNA extracted from stranded *S. chinensis* individuals (i.e., tissue-extracted DNA used for checking intraspecific variation) and synthesised DNAs of the entire *Cytb* region (1,140 bp; Table S3) of the three potentially co-occurring cetacean species: *T. truncatus*, *D. capensis*, and *S. coeruleoalba*. The concentrations of the synthesised DNAs of these three species prepared at 100 and 1,000 copies/µL, which we consider to be the highest expected eDNA concentrations in natural water samples.

After verifying the specificity of the primers and probe, we determined the LOD and LOQ. Different definitions of LOD and LOQ exist, and we followed the definition by Klymus et al. (2020) [38]; in our study, LOD is defined as the lowest concentration of template DNA that produced at least 95% positive detections among replicates, while LOQ is defined as the lowest concentration that could be quantified with a coefficient of variation (C.V.) for DNA copy estimates below 35%. C.V. values were calculated based on C_t_ (the number of PCR cycles that exceed a defined threshold of the fluorescent intensity) and PCR efficiency assuming the log-normal distribution, according to Forootan et al. [39] and Klymus et al. [38], as 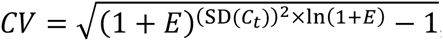, where *E* is the PCR efficiency and SD(*C_t_*) is the standard deviation of replicate C_t_ values. To rigorously determine the LOD, at least 20 replicates are required; we analysed 24 replicates of each standard DNA concentration at 0.5, 1, 10^1^, 10^2^, 10^3^, 10^4^, 10^5^, and 10^6^ copies/µL (equivalent to 1, 2, 2.0× 10^1^, 2.0× 10^2^, 2.0× 10^3^, 2.0× 10^4^, 2.0× 10^5^, 2.0 × 10^6^ copies/reaction, as we used 2 µL DNA for the qPCR analysis) using the protocol described above. The qPCR results were analysed using a custom script provided in Klymus et al. [38]. Briefly, the relationship between standard DNA concentrations and detection probability was modelled using multiple regression models, including asymptotic regression, log-logistic, Michaelis-Menten, Weibull type I, and Weibull type II. Similarly, the relationship between standard DNA concentrations and C.V. values of C_t_ was modelled using exponential decay model, linear model, and polynomial models. The best LOD and LOQ models were chosen to determine “modelled” LOD and LOQ, as explained in the Result section. Detailed methods are described in Klymus et al. [38] and in the custom R script available on their GitHub page (https://github.com/cmerkes/qPCR_LOD_Calc).

### Field survey in Hong Kong waters

To evaluate the performance of the primers and probe on natural water samples, we first collected “proof-of-concept” field samples. On 16 July 2024, we collected two seawater samples (500 and 1,500 mL) from areas where *S. chinensis* were sighted (22°11′32″N, 113°50′57″E; Fig. 1b, indicated by the white diamond). The distance between the sighted *S. chinensis* individuals and the sampling locations was approximately 30 m, leading us to expect detectable amounts of *S. chinensis* eDNA in the seawater samples. To ensure contamination control, we also filtered Milli-Q water as a field negative control. The collected water samples and Milli-Q water were filtered using Sterivex filter cartridges (φ0.45-µm, SVHV010RS; Merck Millipore, Darmstadt, Germany) using a plastic syringe. The samples were stored in a cooler box to preserve eDNA before being transferred to a – 20°C freezer in the laboratory.

Following the analysis of the proof-of-concept field samples, we conducted field surveys in July 2024 in the western and southern Lantau regions. The summer survey took place on 29 July 2024, during which 1 L of surface and bottom water samples were collected from 14 locations. Surface water was collected using a bucket, while bottom water was collected with a Niskin water sampler. The collected water samples were filtered using Sterivex filter cartridges (φ0.45-µm, SVHV010RS) and the gravity filtration method [40] (Fig. 1c). Then, 2 mL of RNAlater was added to preserve eDNA on the boat, and they were transported to a –20°C freezer in the laboratory within several hours after the sample collection. To ensure contamination control, we also collected three field negative control samples (i.e., filtering 1 L of Milli-Q water for each) during the field surveys. The first, second, and third field negative controls were processed at the initial (sP01) and middle (sP04) sampling locations, and immediately after the completion of all sampling, respectively. In total, we collected 31 samples (2 water layers × 14 locations + 3 field negative controls). Water temperature, pressure, salinity, and dissolved oxygen (DO) at the sampling locations were measured using a CTD sensor (Model ASTD102; JFE Advantech Co., Ltd., Nishinomiya, Hyogo, Japan). We sighted several *S. chinensis* individuals during our survey, and the sighting locations were recorded.

### eDNA extraction and TaqMan qPCR analysis

eDNA was extracted from the Sterivex filter cartridges using a protocol described in a previous study [41] with some modifications. First, RNAlater solution (Thermo Fisher Scientific) was removed from the filter cartridge using a vacuum pump. One ml of H_2_O was added to wash the filter cartridge, which was removed using a vacuum pump. Then, Buffer ATL (380 µL) and Proteinase K solution (20 µL) were mixed, and the mixture was added to each filter cartridge. The materials on the cartridge filters were subjected to cell lysis by incubating the filters at 56°C for 30 min. After the incubation, Buffer AL (400 µL) was added to the filter cartridge, and further incubated at 56°C for 10 min to dissolve white precipitation. The incubated and lysed mixture was transferred into a new 2-ml tube from the inlet of the filter cartridge using a manual centrifuge (Handzentrifuge, Hittich, Westphalia, Germany). The collected DNA was purified using a DNeasy Blood & Tissue kit following the manufacturer’s protocol. After the purification, DNA was eluted using Buffer AE (100 µL). Eluted DNA samples were stored at −20°C until further processing.

qPCR analysis was performed following the protocol described in the previous section (i.e., 55 cycles of qPCR using Environmental Master Mix 2.0) with the standard DNAs at the concentrations of 1, 10^1^, 10^2^, 10^3^, and 10^4^ copies/µL. qPCR negative controls were included in the qPCR analysis by replacing the template DNA with ultrapure water. For all samples, qPCR was conducted in three technical replicates, and the average value of the replicates was calculated. “Not detected” was replaced with “0 copies/µL” for all measurements. For the proof-of-concept field samples (i.e., the distance between the sampling locations and the sighted *S. chinensis* individuals was approximately 30 m), we sent the amplified product to a company for Sanger Sequencing to verify if it is *S. chinensis* sequence. Note that specific tests for DNA extraction loss (e.g., spike-in experiments) and PCR amplification inhibition (e.g., internal positive controls) were not conducted in this study.

### Statistical analysis and data visualization

All statistical analyses were performed using R [42]. The LOD and LOQ were determined using the custom R script provided by Klymus et al. [38], as described in the previous section. The qPCR detection was visualized using the ggplot2 [43], ggmap [44], and rphylopic [45] packages of R. Differences in detection rates between surface and bottom water layers were analyzed using McNemar’s test. The relationships between environmental variables and *S. chinensis* eDNA detection were analysed using a generalized linear mixed model (GLMM) with sampling location as a random factor, assuming a binomial distribution. All the code and qPCR data was deposited on Github (https://github.com/ong8181/qpcr-cwd).

## Results

### Validation of qPCR primers and probe

Our primers and probe successfully amplified the synthesised standard DNA (Fig. S1a). Tissue-extracted DNA was also amplified, with target region concentrations estimated at approximately 10^6^–10^7^ copies/µL (Fig. S1b). On the other hand, the synthesised DNA fragments of the three cetacean species (*T. truncatus*, *D. capensis*, and *S. coeruleoalba*) were mostly unamplified, despite the relatively high concentrations of the input DNA (100–1,000 copies/µL) (Fig. 2). Slight amplification was observed in *T. truncatus* (100 copies/µL; Fig. 2f) and *D. capensis* (1,000 copies/µL; Fig. 2e) samples; however, these amplification curves appeared after 40 cycles and negligible, and the resulting signals were far weaker than that of 1 copy/µL of *Sousa chinensis* DNA. These results indicate that the primers and probe are highly specific to *S. chinensis*.

**Figure 2.**
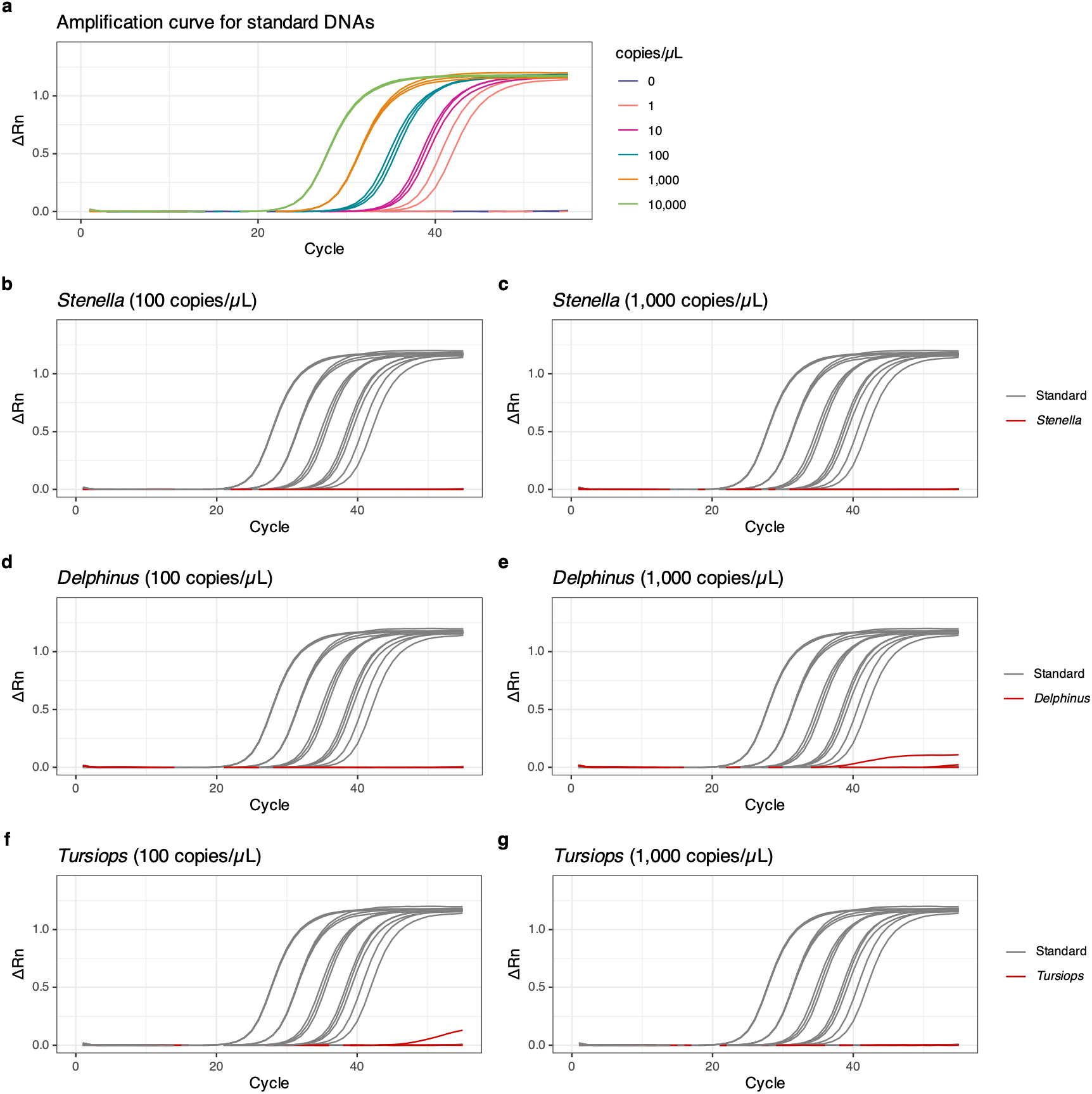
Specificity tests of the *Sousa chinensis* qPCR assay against three sympatric Delphinidae species. **(a)** Amplification curves of S. *chinensis* standard DNA across a dilution series (0 to 10,000 copies/ µL). Different colors indicate different template concentrations. **(b-g)** Amplification curves of syn-thesized target-region (Cytb) DNA from three non-target, but potentially co-occurring cetacean species (red curves) tested at 100 copies/ µL **(b, d, f;** slightly exceeding the maximum concentration observed in our "proof-of-concept" field samples, -60 copies/ µL) and 1,000 copies/ µL **(c, e, g),** plotted alongside the S. *chi-nensis* standard DNA curves (gray curves, 0 to 10,000 copies/ µL): **(b, c)** *Stenella coeruleoalba,* **(d, e)** *Delphinus capensis* and (f, g) *Tursiops truncatus.*

The LOD and LOQ were determined by the qPCR analysis of 24 replicates of the standard DNA of 1–2.0 × 10^6^ copies/reaction (0.5–10^6^ copies/µL) (Fig. 3a and Tables S5– S6). The R^2^ of the standard curve was 0.998 and PCR efficiency was 100.1%. The discrete LOD was 20 copies/reaction, a concentration at which all 24 replicates were successfully detected (Table S6). The best fit model for the LOD was Weibull type II, and the LOD was 3.22 copies/reaction (95% confidence interval [CI] is 2.01–4.45 copies/reaction), equivalent to 1.61 copies/µL (95% CI is 1.00–2.22 copies/µL) (Fig. 3b). Other models produced similar modelled LODs, ranging from 3.22 to 4.29 copies/reaction (Table S7).

**Figure 3.**
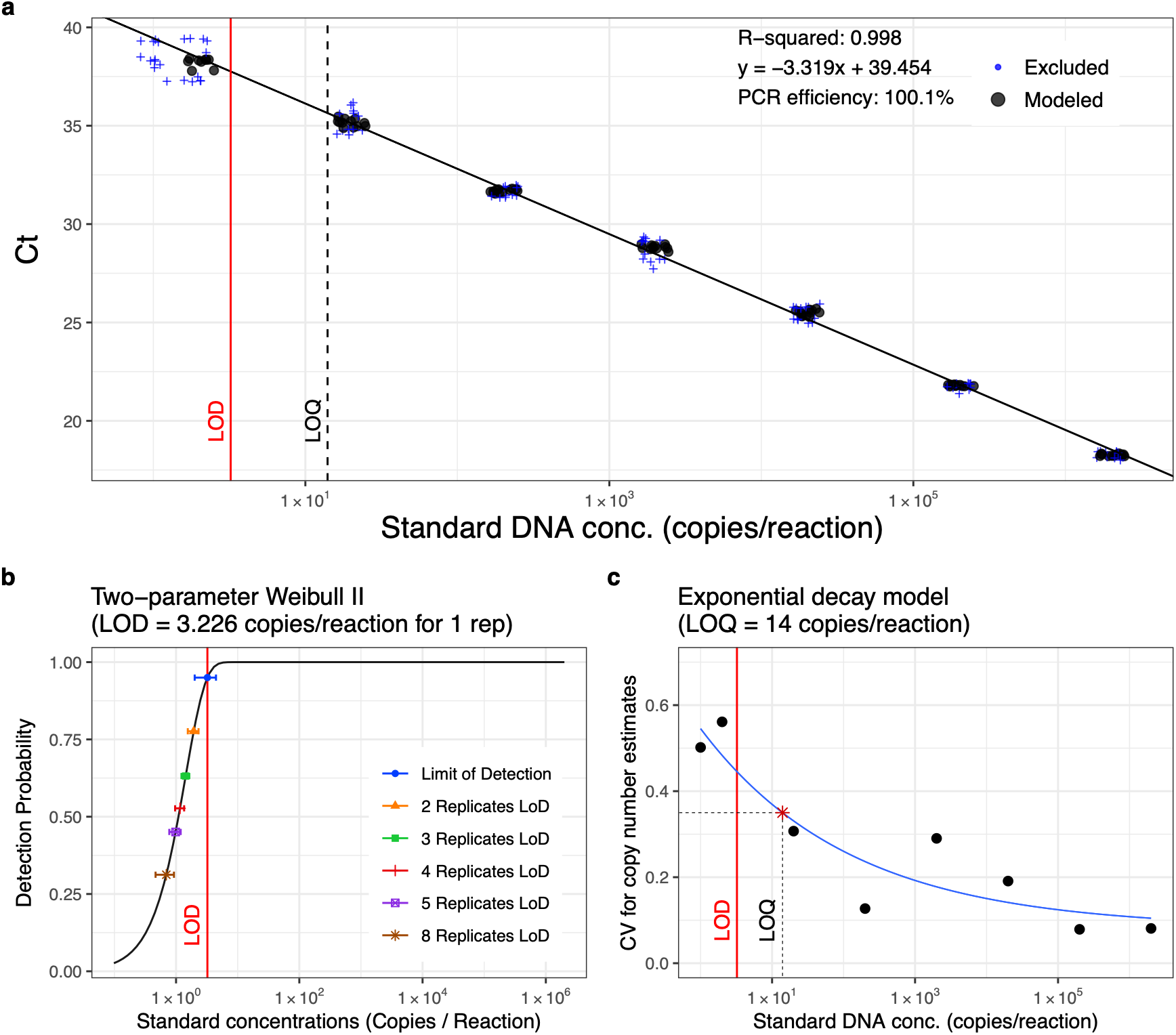
Standard curve, the limit of detection (LOD) and limit of quantification (LOQ) of the primers and probe specific to *Sousa chinensis.* **(a)** Standard curve generated from 24 technical replicates across eight standard DNA concentrations (1 to 2.0 × 10^6^ copies/ reaction). The x-axis represents copy numbers of the *Cytb* target region per qPCR reaction, and the y-axis represents the *C_t_* value. Following Klymus et al. (2020), filled black points indicate data included in the linear regression (the middle two quartiles, i.e., 25th-75th percentiles, of standards with ≥ 50% detection rates), whereas blue "+" symbols indicate data excluded from the regression (points outside the middle two quartiles or standards with < 50% detection rates, i.e., 1 copy/ reaction). The red solid vertical line and black dashed vertical line indicate the modeled LOD (3.22 copies/ reaction) and modeled LOQ (14 copies/ reaction), respectively. (b) Determination of the LOD. Open circles indicate observed detection rates at each standard concentration. The two-parameter Weibull type II model (W2. 2 () in R) was selected as the best-fitting model. Colored symbols with horizontal error bars indicate effective LODs (with 95% confidence intervals) for different numbers of technical replicates (1 to 8 replicates). (c) Determination of the LOQ based on the exponential decay model. The x-axis represents copy numbers and the *y*-axis represents the coefficient of variation (CV) for copy number estimates calculated following Forootan et al. (2017) and Klymus et al. (2020). The blue curve indicates the exponential decay model (see Fig. S2 and Table S7 for comparisons of all candidate models). The red vertical line indicates the modeled LOD, and the black "*" symbol indicates the 35% CV precision threshold, at which the copy number is defined as the modeled LOQ (14 copies/reaction, equivalent to 7.0 copies/ *μ*L).

The discrete LOQ was 20 copies/reaction (Table S6). To determine the modelled LOQ, the relationship between C.V. and standard DNA concentrations was modelled and visualized (Fig. S2). Although more complex models, such as 4^th^-, 5^th^-, and 6^th^-order polynomial models, yielded lower AIC and BIC values (Table S7), we selected a simpler model, the exponential decay model, as the most appropriate, given that only eight standard DNA concentrations were tested (i.e., a small degree of freedom). The LOQ based on the exponential decay model was 14 copies/reaction (7.0 copies/µL) (Fig. 3c). LOQs derived from other models were also similar, ranging from 8 to 24 copies/reaction, except for the linear model (50 copies/reaction), which produced the second-highest AIC and BIC values (Table S7).

### Raw results and data pre-processing for qPCR

All raw data are provided in Tables S8–S12. Table S8 presents the summarized data used for subsequent statistical analyses. Tables S9 (surface sample analysis plate) and S10 (bottom sample analysis plate) contain the analytical metrics, including raw C_t_ values, PCR efficiencies, C_t_ thresholds, and standard curve R^2^ values. Tables S11 and S12 provide the raw amplification curve data; an interactive visualization is also available at https://github.com/ong8181/qpcr-cwd/data_supplementary/. For standard curve calibration, specific outliers (e.g., standard DNAs at low concentrations) were excluded from the construction of the standard curve (see Tables S9 and S10 for details). Additionally, three technical replicates with invalid amplification curves (one replicate each from samples sP01-B, sP02-B, and sP03-B) were excluded from subsequent analyses.

### Proof-of-concept field samples in Hong Kong waters

The concentrations of *S. chinensis* eDNA in the two “proof-of-concept” field samples were 4,063 and 9,136 copies/L seawater (60.95 copies/µL and 45.68 copies/µL in the template DNA, respectively), while no *S. chinensis* eDNA was detected in the field negative control (Table S9). These values exceeded the discrete and modelled LOD and LOQ. In addition, the sequences of the amplified eDNA were confirmed as *S. chinensis* DNA using Sanger sequencing (Table S13). This result suggests that our qPCR primers and probe can detect *S. chinensis* eDNA in the natural coastal ecosystem.

### Field survey in Hong Kong waters

The physicochemical properties of the sampling locations during the survey are shown in Fig. S3. Water temperature was 26.3–28.9°C, salinity was 13.1–29.3‰, and dissolved oxygen was 2.41–6.79 mg/L (Fig. S3a–c). Salinity was relatively low due to freshwater discharge from the Pearl River, and rainfall during the survey further contributed to the low salinity, especially at the surface layers.

Among all 81 replicates in the field samples (three invalid amplification excluded), 22 positive amplifications were observed, and 21 amplifications were above the modelled LOD. Among them, six amplifications were above the modelled LOQ (7.0 copies/µL) (Table S8). However, we detected the mean value of 2.87 copies/µL of *S. chinensis* eDNA in one of the three field negative controls collected at the first sampling location (sP01), which was higher than the modelled LOD (1.61 copies/µL or 3.22 copies/reaction). To ensure conservative analysis and avoid false positives, we set the mean value of 2.87 copies/µL as a conservative positivity threshold (or contamination-adjusted cut-off) for the field samples, rather than relying on the technical (modelled) LOD determined for our qPCR. In addition, a sample was classified as positive only if at least two out of the three technical replicates exceeded the modelled LOD (1.61 copies/µL).

In total, six positive detections among 28 samples were classified as positive (Fig. 4). The mean eDNA concentration of the six positive samples was 5.70 copies/µL, equivalent to 570 copies/L seawater, but only one sample was above the modelled LOQ (the maximum concentration, 10.87 copies/µL, or 1,087 copies/L seawater, was detected in the bottom water sample at sP03). Therefore, only “qualitative” results (detected or not) are reported in Fig. 4. We sighted one *S. chinensis* individual at the location sP02 (the pink silhouette in Fig. 4), and *S. chinensis* eDNA was detected from both surface and bottom water samples at the location. The eDNA concentrations are provided in the supporting information (Fig. S4a, b and Table S8). We found no statistically clear effects of physicochemical properties or water layers on eDNA detection (all statistical tests, *P* > 0.05; Fig. S5).

**Figure 4.**
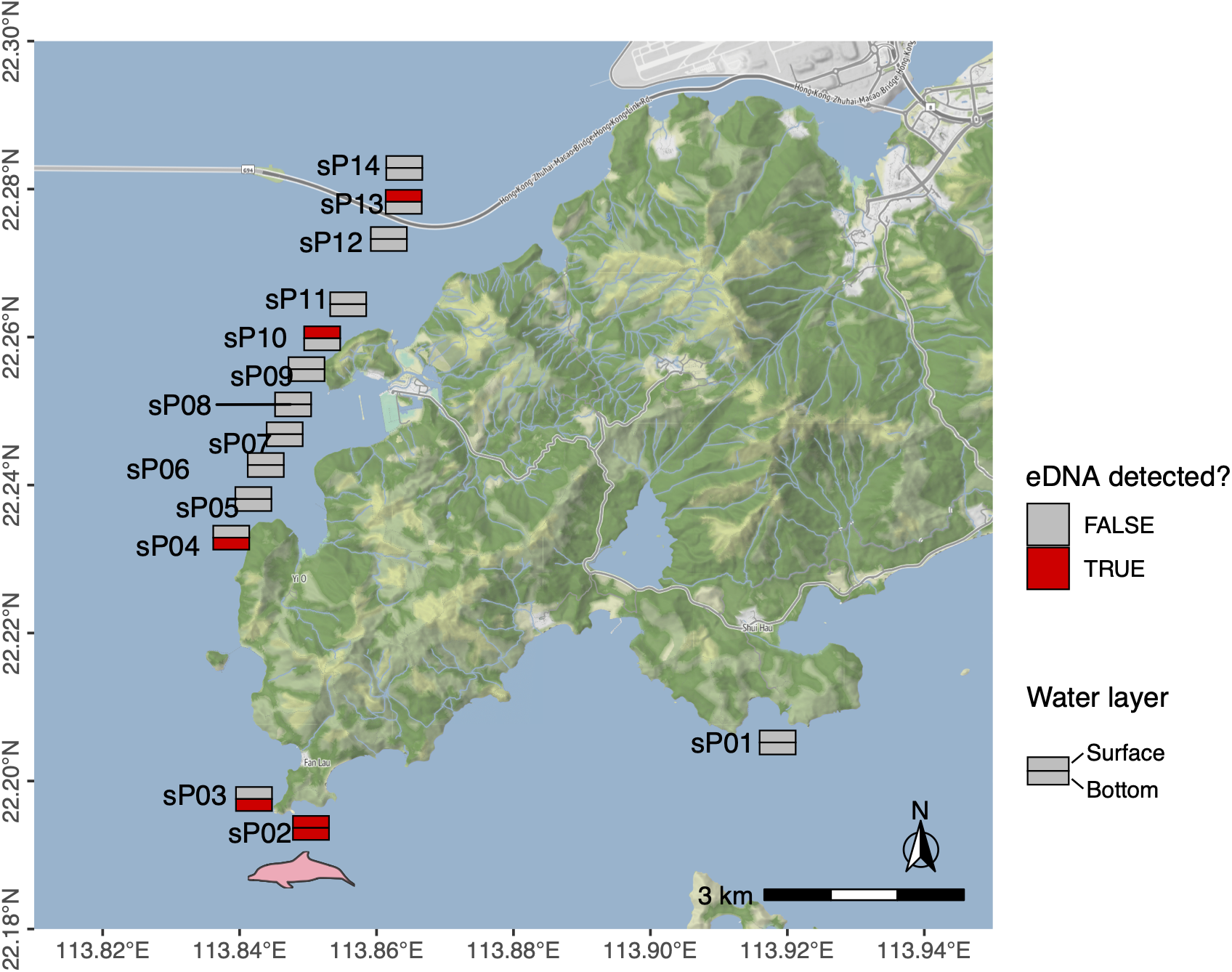
Detection of *Sousa chinensis* eDNA from field seawater samples using qPCR. qPCR-based detection of S. *chinensis* eDNA from surface and bottom water samples taken in July 2024. Labels indicate the sample collection locations (see the metadata in the supporting information). Red and gray color indicate positive and negative detection, respectively. Upper and lower rectangles indicate the result for surface and bottom water layers, respectively. S. *chinensis* silhouette indicates S. *chinensis* sighting during the field survey (silhouette credit, Chris huh, CC BY-SA, https://creativecommons.org/licenses/by-sa/3.0/). Map data by OpenMapTiles and OpenStreetMap contributors, under ODbL.

## Discussion

### Validation of the qPCR primers and probe

*In silico* evaluation using Primer-BLAST and experimental validation showed that our qPCR primers and probe were highly specific to *S. chinensis*. The synthesised DNAs of the three cetacean species (*T. truncatus*, *D. capensis*, and *S. coeruleoalba*), which are occasionally found in Hong Kong waters, were not amplified, despite their relatively high concentrations (100–1,000 copies/µL) in the experiment. Because cetacean eDNA concentrations would be expected to be low in most cases (e.g., below 10^3^–10^4^ copies/L water; [46]), our qPCR primers and probe are expected to amplify only *S. chinensis* eDNA in samples from natural ecosystems.

The discrete and modelled LOD and LOQ we determined were comparable to those reported in Klymus et al. [38]. According to Table 1 in Klymus et al. [38], the mean modelled LOD is 7.45 copies/reaction, with most values below 10 copies/reaction. The mean modelled LOQ is 88.2 copies/reaction, with most values below 50 copies/reaction. In our study, the modelled LOD and LOQ are 3.22 copies/reaction (1.61 copies/µL) and 14 copies/reaction (7.0 copies/µL), respectively. This result, along with the specificity test mentioned above, suggests that our qPCR primers and probe have sufficient sensitivity and specificity to analyse natural seawater samples. Indeed, we detected *S. chinensis* eDNA in the proof-of-concept field samples, and their concentrations were higher than the LOQ (4,063 and 9,136 copies/L seawater, or 60.95 and 45.68 copies/µL), further supporting the performance of our qPCR primers and probe.

### Detection of *Sousa chinensis* eDNA from Hong Kong seawater samples

We successfully detected *S. chinensis* eDNA in the field water samples (Fig. 4), but the eDNA concentrations were below the modelled LOQ in most samples. The mean eDNA concentrations of the six positive samples, although five of them are below the modeld LOQ, were 570 copies/L seawater. Baker et al. (2018) reported an average of 204 copies/L seawater of killer whale eDNA (4.08 copies/µL in 50-µL of extracted DNA from 1 L of seawater) [46], which were similar to the values in our study. Lin and Ushio (2026) also reported similar concentrations for the Indo-Pacific finless porpoise eDNA in the same region (the mean eDNA concentration of positive samples was 322 copies/L seawater) [18]. These low eDNA concentrations may be common for cetacean species. However, when we collected the proof-of-concept field samples approximately 30 m from the *S. chinensis* individual (closer than the distance to *S. chinensis* individuals we sighted during the field survey), we detected eDNA concentrations higher than the modelled LOQ. Therefore, the low eDNA concentrations in our surveys could be because *S. chinensis* individuals were relatively far from the sampling locations (e.g., > 30 m). Alternatively, *S. chinensis* eDNA may have been dispersed in the opposite direction from our sampling location due to sea currents.

Our July 2024 survey showed that *S. chinensis* eDNA was detected in the western Lantau region, but not at the single location in the southern Lantau region (sP01) (Fig. 4). The western Lantau detections align with AFCD monitoring data showing high utilization of this area by *S. chinensis* [47] (Fig. 1b), whereas broader surveys are needed to evaluate their presence in southern Lantau, an important habitat for another resident cetacean species, Indo-Pacific finless porpoises (*N. phocaenoides*) [26]. While positive detections at the sighting station (sP02) and in our near-dolphin proof-of-concept samples (Fig. 1b) suggest that proximity to dolphins enhances detection probability, detections at locations without sightings could reflect either visually cryptic (e.g., submerged) individuals nearby or passive eDNA transport via estuarine currents. However, our study was not designed to provide the direct evidence; the importance of the distance in detecting *S. chinensis* eDNA and the capability of eDNA to detect visually undetectable individuals should be quantitatively evaluated in future studies. Additionally, because our single-season pilot survey lacked independent field replicates, hydrodynamic modeling, and PCR inhibition tests, future studies incorporating these elements are required to disentangle local presence from eDNA transport and false negatives.

Our analysis did not find any statistically clear influences of physicochemical properties and water layers on eDNA detection (Fig. S5). While there were differences in physicochemical properties between surface and bottom water (Fig. S3) and some fish eDNA studies reported that different species were detected in the surface and bottom water layers [48,49], overall detection rates were identical between surface (3/14) and bottom (3/14) waters (Figs. 4 and S5a). However, statistical power was limited by our small sample size (14 locations). Thus, the lack of a statistically clear difference may not imply equivalence across depths.

### Future perspectives and implications for conservation

To implement more effective *S. chinensis* monitoring using qPCR-based eDNA analysis, future studies may consider the following improvements. First, increasing the volume of water samples can enhance detection probability. In theory, a larger water volume will capture more eDNA on the filter, increasing the probability of positive amplification in qPCR. Indeed, How et al. (2024) demonstrated that in Hong Kong waters, greater water volume leads to an increased number of detected species and reduced variation among technical replicates [50]. They recommended filtering ≥ 2 L of water samples for better detection. Yoshida et al. (2023) also evaluated the effect of water volume on the number of detected species, suggesting that 10–20 L of seawater may be suitable for deep-sea fish eDNA detection [51]. While the optimal water volume varies depending on the target species and ecosystem, processing larger volumes of water than we did in our study would likely increase the detection probability for small cetaceans in Hong Kong waters, provided that the associated time- and labour-costs are acceptable.

Second, since other cetacean species, including the Indo-Pacific finless porpoise (*N. phocaenoides*), may be present or appear in Hong Kong waters, applying eDNA metabarcoding would be beneficial for detecting multiple cetacean species at once. Ushio et al. (2025) have developed cetacean-specific eDNA metabarcoding primers, named µCeta [17], which would improve the efficiency of cetacean monitoring in Hong Kong waters.

Recently, Lin and Ushio (2026) applied µCeta eDNA metabarcoding in the Soko Islands region of southern Lantau and detected *N. phocaenoides* and the false killer whale (*Pseudorca crassidens*), a transient cetacean species in Hong Kong waters [18], demonstrating the advantage of eDNA metabarcoding for detecting multiple cetacean species simultaneously.

Third, this eDNA metabarcoding approach could be expanded to assess prey communities (i.e., fish eDNA metabarcoding). Analysing co-occurrence patterns of cetaceans and fish would help us understand the factors influencing the distribution patterns of *S. chinensis*, which will contribute to more effective conservation of *S. chinensis*. Although eDNA metabarcoding generally involves a tradeoff of decreased detection sensitivity for a single target species compared to species-specific qPCR approach, the comprehensive ecological insights provided by this “multi-taxa” approach may outweigh the sensitivity limitations.

Lastly, every survey method has its own advantages and limitations. As demonstrated in this study, *S. chinensis* eDNA concentrations can be low, potentially limiting its detection sensitivity compared to other methods depending on environmental conditions. While water sampling requires no specialized skills, laboratory and sequencing expenses may make eDNA analysis more costly than single visual surveys. Furthermore, currently, eDNA alone cannot provide demographic data such as individual size, age, or sex. However, eDNA analysis complements traditional surveys, e.g., by detecting intra-specific variations [52] and underwater individuals. Additionally, as eDNA technologies are rapidly being developed, it may become possible to estimate physiological status using eDNA (e.g., by analysing DNA methylation patterns) [53]. Therefore, selecting or combining appropriate methods based on available financial, human, and technological resources is crucial for more effective biodiversity monitoring.

## Conclusions

In the present study, we rigorously designed and validated species-specific qPCR primers and probe for detecting *S. chinensis*. By confirming its high sensitivity and specificity against sympatric cetaceans, we have added a robust and reliable tool to the expanding suite of eDNA markers for marine life. Furthermore, our pilot survey in Hong Kong waters successfully demonstrated the practical applicability of this assay in natural estuarine environments. Looking forward, with further improvements to the eDNA-based *S. chinensis* survey method, such as increasing water volume, expanding spatial coverage, and targeting other cetaceans and fish, our eDNA-based cetacean monitoring framework will contribute to more effective conservation of *S. chinensis* in Hong Kong waters.

## Data and Code Accessibility

All scripts and raw data except for the sequence data used in the present study are available on Github (https://github.com/ong8181/qpcr-cwd). Sanger sequence data was deposited in DDBJ Nucleotide Sequence Submission System (NSSS) (Accession No = LC899610–LC899638).

## Declaration of AI use

We used generative AIs to improve the clarity of the manuscript text. We reviewed and edited all AI-generated content to ensure accuracy and appropriateness.

## Acknowledgments

We thank Suixuan Huang, Ming-Wai Li, Lucia Hu, and Chengbin Liu for their assistance in the field work and Leung Kwan Chak for his assistance in the tissue sample collection and management. We thank the Agriculture, Fisheries and Conservation Department (AFCD) of Hong Kong, and the Ocean Park Hong Kong (OPHK) and Ocean Park Conservation Foundation Hong Kong (OPCFHK) for their permission for using cetacean tissue samples and assistance in collecting them.

## Author contributions

MU conceived and designed research; ROK, ST, and MU designed primers; ROK and MU validated primers; ROK, TO, XL, LP, EM, and MU performed field water sampling; ROK and XL performed qPCR experiments with technical advice from ST and MU; ROK, TO, and MU analysed the data; ROK and MU wrote the first draft; MU completed the first draft; MU revised the manuscript and all authors discussed the results and completed the manuscript.

## Supporting information files

Supplementary Tables.xlsx

Table S1. List of mammal species for primer design

Table S2. Metadata of stranded individuals used for Sanger sequencing to assess intraspecific variation

Table S3. Summary for the primer-BLAST results Table S4. Synthesized DNA sequences

Table S5. Raw data for amplification curve for standard DNA

Table S6. Ct values and summary statistics for LOD and LOQ analysis Table S7. LOD and LOQ model summary

Table S8. Summary table for the qPCR results for field samples Table S9. qPCR results for Plate 1 (surface samples)

Table S10. qPCR results for Plate 2 (bottom samples)

Table S11. Raw data for amplification curve for Plate 1 (surface samples)

Table S12. Raw data for amplification curve for Plate 2 (bottom samples) Table S13. Sanger sequence data for "field positive" samples

Supplementary Figures.pdf

Figure S1. Amplification of *Sousa chinensis* standard DNA and tissue-extracted DNA

Figure S2. Comparison of candidate models for determining the limit of quantification (LOQ) of the *Sousa chinensis* qPCR assay.

Figure S3. Physical profiles of seawater at the sampling locations.

Figure S4. Concentrations of *Sousa chinensis* eDNA from field seawater samples using qPCR.

Figure S5. The relationship between seawater physical properties and qPCR detection in July 2024.

