## Supplementary Figures for "Detection of environmental DNA of the Indo-Pacific humpback dolphins in Hong Kong waters using quantitative PCR"

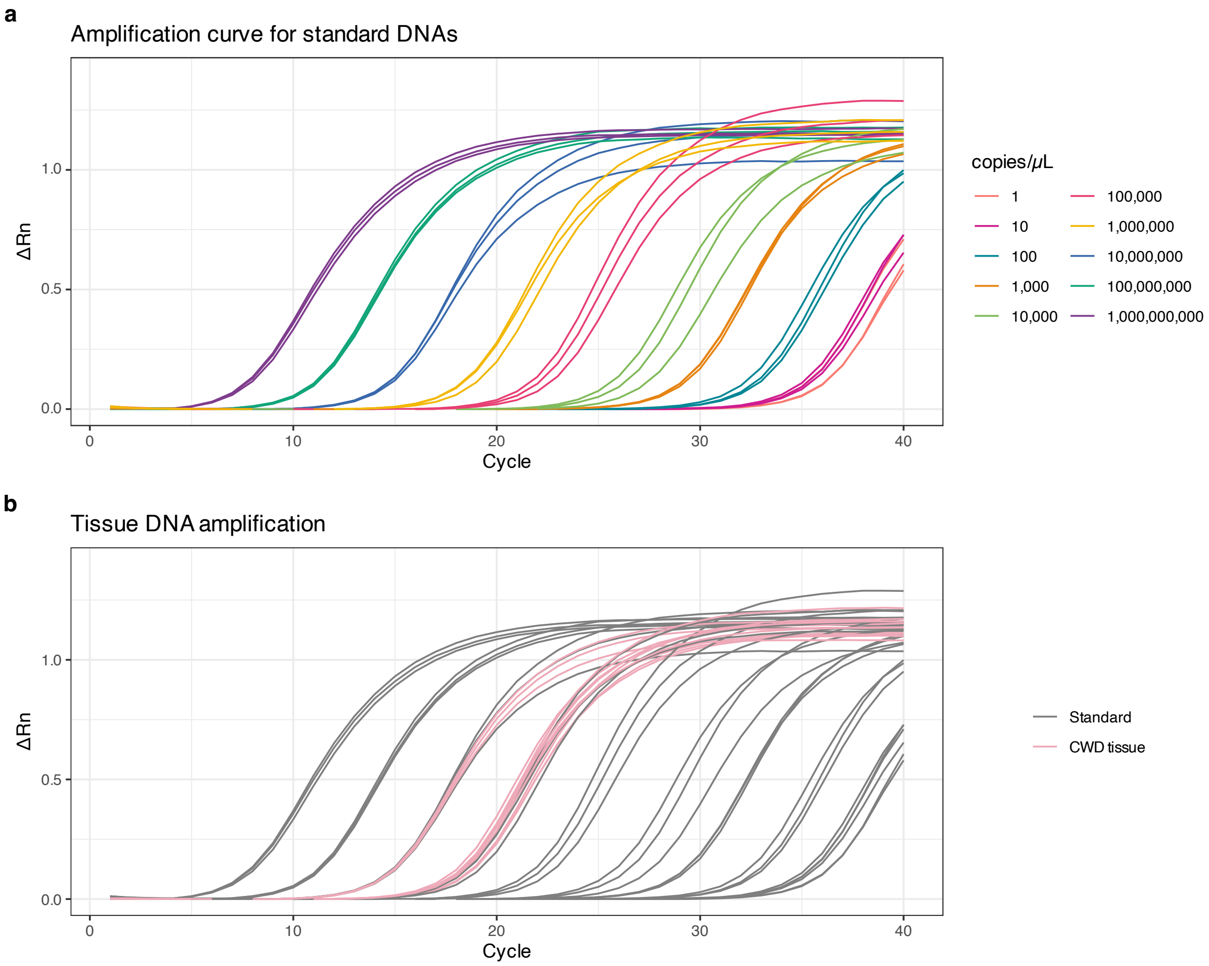


**Figure S1 | Amplification of *Sousa chinensis* standard DNA and tissue-extracted DNA.** (**a**) Amplification curve of the target region (*Cytb*) of *Sousa chinensis* (synthesized DNA). Different colors indicate different concentrations. Amplification curve of (**a**) S. chinensis standard DNAs and (**b**) *S. chinensis* tissue DNA (pink curve). Gray curves in **b** indicate amplification curves of different concentrations of *S. chinensis* standard DNA.

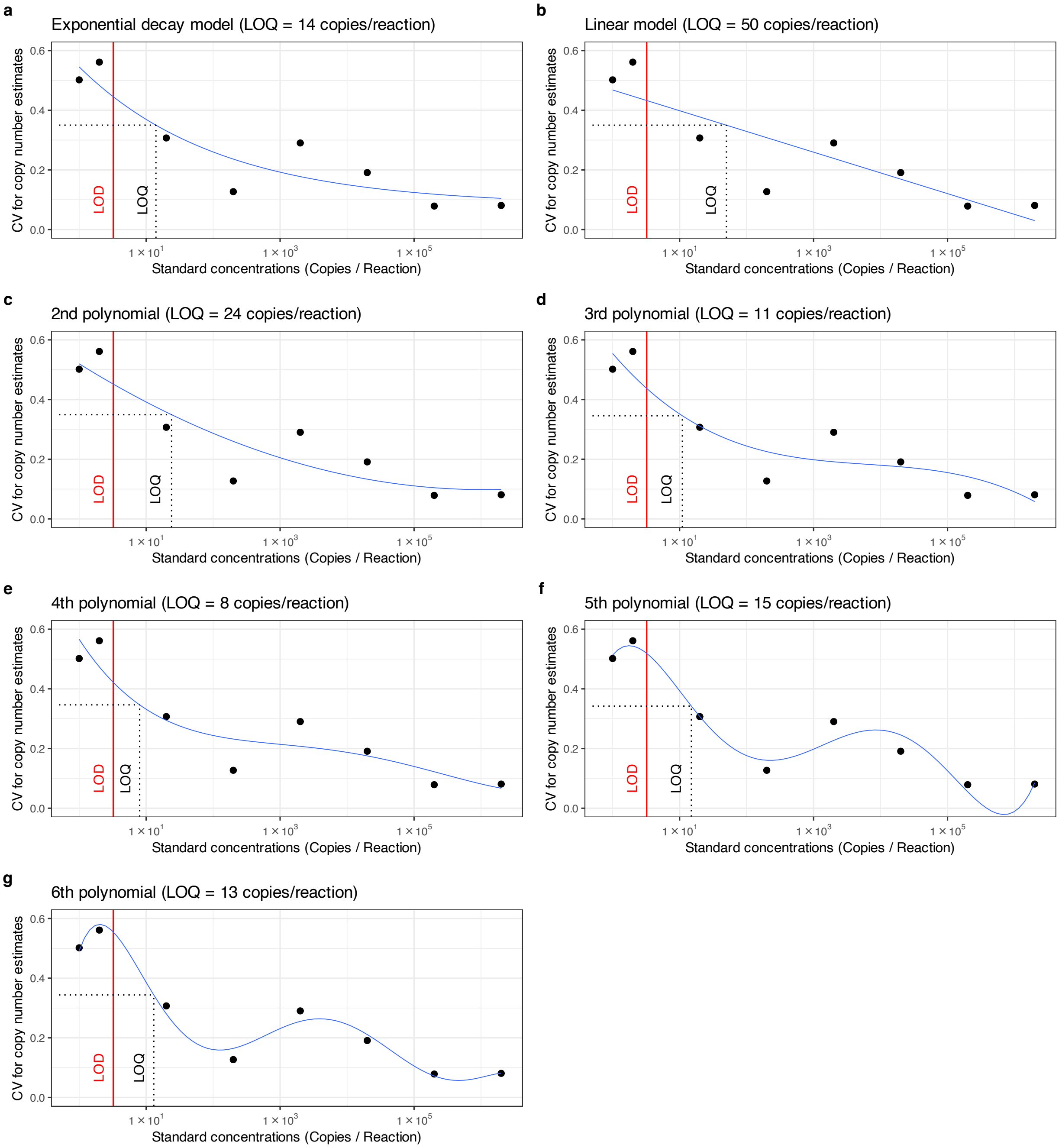


**Figure S2 | Comparison of candidate models for determining the limit of quantification (LOQ) of the *Sousa chinensis* qPCR assay.** (**a**) Exponential decay, (**b**) linear, and (**c–g**) 2nd- to 6th-order polynomial models. The $x$-axis represents standard DNA concentrations (copies/reaction), and the $y$-axis represents the coefficient of variation (CV) for copy number estimates calculated following Forootan et al. (2017) and Klymus et al. (2020). Black points indicate observed CVs at each standard concentration ($n = 24$ replicates), and blue curves indicate the fitted models. Red vertical lines indicate the modeled LOD (3.22 copies/reaction), and black dotted lines indicate the 35% CV threshold (horizontal) and the resulting modeled LOQ (vertical). Model diagnostics (RSE, AIC, AICc, and BIC) for all seven models are provided in Table S7.

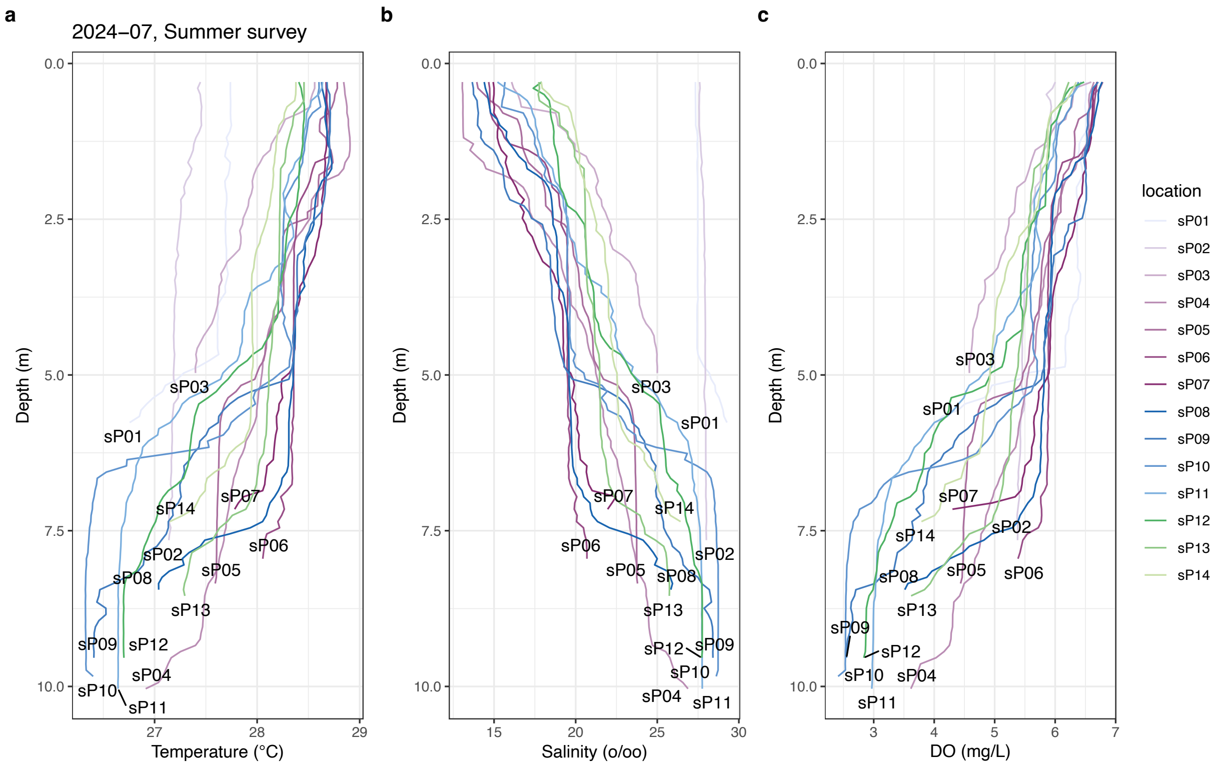


**Figure S3 | Physical profiles of seawater at the sampling locations.** (**a**) Temperature, (**b**) salinity, and (**c**) dissolved oxygen (DO) at each sampling location for the summer survey in July 2024. Different colors indicate different sampling locations, and labels indicate sampling locations.

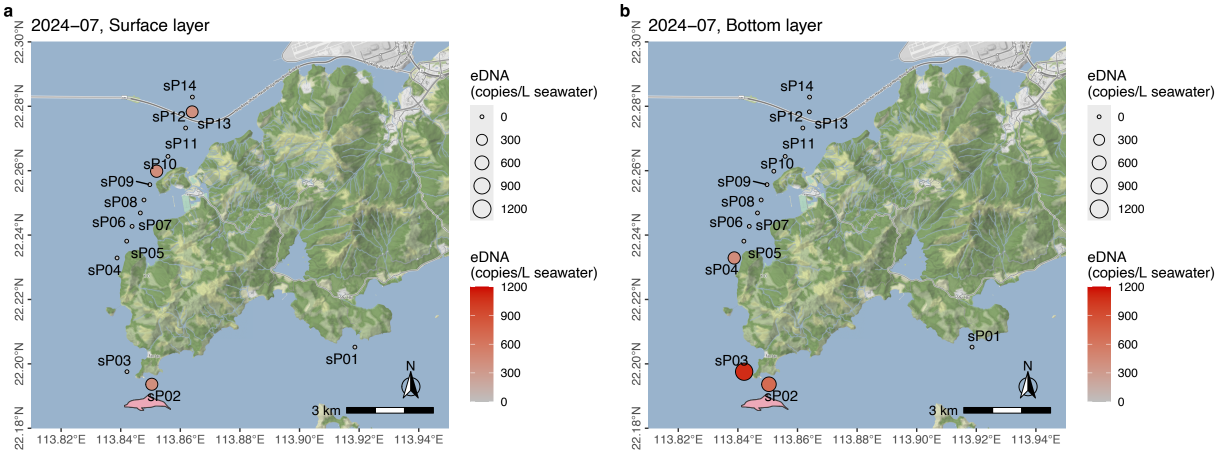


**Figure S4 | Concentrations of *Sousa chinensis* eDNA from field seawater samples using qPCR.** Concentrations of *S. chinensis* eDNA from (**a**) surface and (**b**) bottom water samples taken in July 2024. Labels indicate sample collection locations. Color density and size of each point represent eDNA concentration. Most the eDNA concentrations are below the limit of quantification (LOQ: 14 copies/reaction or 7 copies/$\mu$L) and are therefore unreliable, but the results are provided as supplementary information. *S. chinensis* silhouette indicates *S. chinensis* sighting during the field survey (silhouette credit, Chris huh, CC BY-SA, https://creativecommons.org/licenses/by-sa/3.0/). Map tiles by Stamen Design, hosted by Stadia Maps, under CC BY 4.0. Map data by OpenMapTiles and OpenStreetMap contributors, under ODbL.

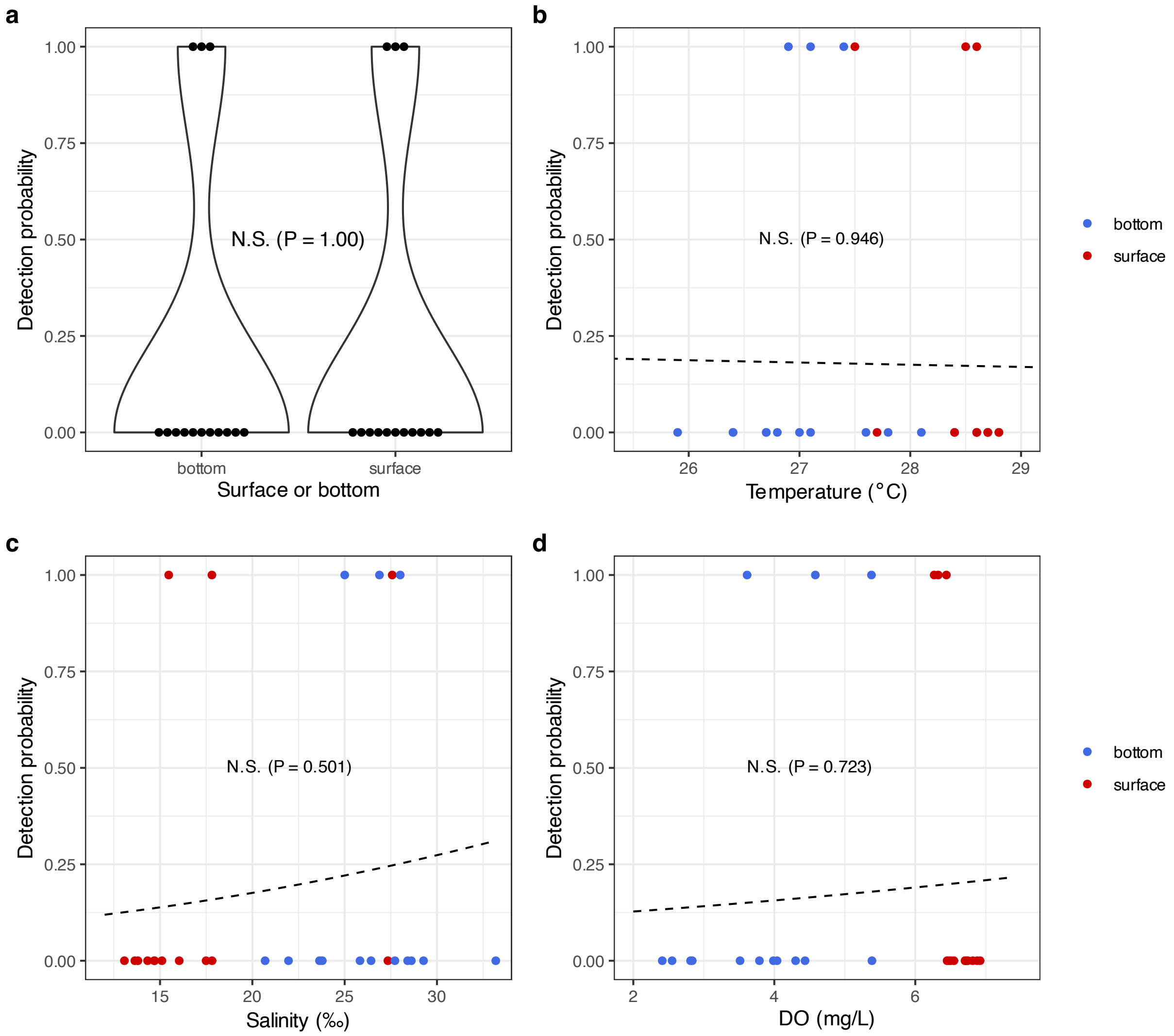


**Figure S5 | The relationship between seawater physical properties and qPCR detection in July 2024.** The relationship between qPCR detection of *Sousa chinensis* eDNA and (**a**) the origin of water samples (surface or bottom), (**b**) temperature, (**c**) salinity, and (**d**) dissolved oxygen (DO), respectively. Red and blue colors indicate surface and bottom water samples, respectively. Lines in **b**-**d** indicate model predictions based on binomial GLMM with each sampling location as a random factor (`glmer(detection ~ environmental variable + (1 | location), family = binomial)`), although all the regressions are not statistically significant. Note that only qualitative results were used for this analysis because the eDNA concentrations of most samples were below the limit of quantification (LOQ).
